# Large-scale comparative small RNA analyses reveal genomic structural variants in driving expression dynamics and differential selection pressures on distinct small RNA classes during tomato domestication

**DOI:** 10.1101/2021.09.25.461803

**Authors:** You Qing, Yi Zheng, Sizolwenkosi Mlotshwa, Heather N. Smith, Xin Wang, Xuyang Zhai, Esther van der Knaap, Ying Wang, Zhangjun Fei

**Affiliations:** Beijing Key Laboratory for Agricultural Application and New Technique, College of Plant Science and Technology, Beijing University of Agriculture, Beijing, China 102206; Bioinformatics Center, Beijing University of Agriculture, Beijing, China 102206; Boyce Thompson Institute, Cornell University, Ithaca, NY 14853; Department of Molecular Genetics, Ohio State University, Columbus, OH 43210; Department of Biological Sciences, Mississippi State University, Starkville, MS 39759; Center for Applied Genetic Technologies, University of Georgia, Athens, GA, 30602; Institute for Plant Breeding, Genetics and Genomics, University of Georgia, Athens, GA, 30602; Department of Horticulture, University of Georgia, Athens, GA, 30602; USDA-ARS, Robert W. Holley Center for Agriculture and Health, Ithaca, NY, 14853; Department of Plant Pathology, Ohio State University, Wooster, OH 44691; Department of Biological Sciences, Louisiana State University, Baton Rouge, LA 70803

## Abstract

Tomato has undergone extensive selections during domestication. Recent progress has shown that genomic structural variants (SVs) have contributed to gene expression dynamics during tomato domestication, resulting in changes of important traits. Here, through comprehensive analyses of small RNAs (sRNAs) from nine representative tomato accessions, we demonstrate that SVs substantially contribute to the dynamic expression of the three major classes of plant sRNAs: microRNAs (miRNAs), phased secondary short interfering RNAs (phasiRNAs), and 24-nt heterochromatic siRNAs (hc-siRNAs). Changes in the abundance of phasiRNAs and 24-nt hc-siRNAs likely contribute to the alteration of mRNA gene expression during tomato’s recent evolution, particularly for genes associated with biotic and abiotic stress tolerance. We also observe that miRNA expression dynamics are associated with imprecise processing, alternative miRNA-miRNA* selections, and SVs. SVs mainly affect the expression of less-conserved miRNAs that do not have established regulatory functions or low abundant members in highly expressed miRNA families, highlighting different selection pressures on miRNAs compared to phasiRNAs and 24-nt hc-siRNAs. Our findings provide insights into plant sRNA evolution as well as SV-based gene regulation during crop domestication. Furthermore, our dataset provides a rich resource for mining the sRNA regulatory network in tomato.

## Introduction

Tomato is the world leading fruit crop in terms of total production and market value (http://www.fao.org/faostat). Originally domesticated in Northern Ecuador and Peru, tomato underwent further selections in Central America and Mexico prior to its arrival in Europe in the early 16^th^ century [1-3]. Along the way, selections had been made for larger fruit, enhanced flavor, and improved resistance to biotic and abiotic stresses [4]. These phenotypic changes reflect the alterations in gene sequences and expression.

Recent evidence has demonstrated that genomic structural variants (SVs) are strongly associated with selection pressure over the course of tomato’s recent evolution that impact the expression of genes underlying certain agronomic traits [5, 6]. SVs include insertions, deletions, duplications, inversions and translocations, and many of them serve as the causative genetic variants for diverse crop traits that have been selected during domestication [5, 6]. For example, the decrease of fruit lycopene levels is strongly associated with deletions in the promoters of multiple key genes involved in lycopene biogenesis in modern tomato [5]. However, the molecular basis underlying the link between genome-wide SVs and gene expression often remains elusive.

Small RNA (sRNA)-mediated gene silencing acts as a key mechanism in regulating gene expression in most eukaryotic organisms. In plants, there are three major groups of sRNAs: microRNAs (miRNAs), phased secondary short interfering RNAs (phasiRNAs), and 24-nt heterochromatic siRNAs (hc-siRNAs) [7, 8]. MiRNAs, phasiRNAs and hc-siRNAs are generated by Dicer-like enzymes (DCLs): DCL1, DCL4, and DCL3, respectively [9]. After production, they are loaded into the RNA-induced silencing complex (RISC) for function [10]. In plants, miRNAs and phasiRNAs with a length of 21 or 22 nt mainly guide cleavage of target mRNAs. By contrast, 24-nt hc-siRNAs play a major role in RNA-directed DNA methylation to confer epigenetic regulation over gene expression [7]. It is known that sRNA-based regulation relies on the abundance of the sRNAs [11]; therefore sRNA abundance has a significant impact on the functions.

In plants, miRNAs and phasiRNAs among distinct species are evolutionarily fluid [12, 13]. Comparative studies on miRNA gene evolution in *Arabidopsis lyrata* and *A. thaliana* that diverged more than 10 million years ago have discovered numerous less conserved miRNA genes that exhibit high divergence in hairpin structures, processing fidelity, and target complementarity [14, 15]. With the increasing number of analyses on sRNA sequencing (sRNA-Seq) data, more and more less conserved miRNAs and phasiRNA-generating loci (PHASs) have been uncovered in diverse species from green algae to flowering plants [12, 13, 16, 17]. However, whether and how the expression patterns and functions of sRNAs have been changed in shorter evolutionary times such as during crop domestication is unknown. Tomato evolved from a wild red-fruited progenitor species, *Solanum pimpinellifolium* (SP) approximately 80 thousand years ago into *S. lycopersicum* var. *cerasiforme* (SLC) [3, 18]. Semi-domesticated SLC further evolved into the fully domesticated tomato, *S. lycopersicum* var. *lycopersicum* (SLL). The evolution within the red-fruited tomato clade provides a unique system for studying selection and functional divergence of plant sRNAs in a shorter time scale thanks to its extensive genetic and genomic resources. Analysis of samples from wild and domesticated tomato accessions may reveal novel regulatory details underlying sRNA evolution and function.

Here, we present comprehensive sRNA profiles of cultivated tomatoes and their wild progenitors, and the novel discovery that genomic SVs can substantially influence sRNA expression dynamics. Our findings show that SVs are an important driving force for the dynamic expression of sRNAs. Moreover, we show that SVs can change the hc-siRNA hotspots in promoters of nearly 100 protein-coding genes, thereby altering their expression. These genes are mostly associated with responses to biotic and abiotic stresses. SVs are also correlated with many rapid birth and death of PHASs that are overwhelmingly related to disease resistance traits. SVs overlapping with miRNA genes can determine the gain or loss of certain less-conserved miRNA genes or affect the expression of miRNAs. Interestingly, the differential expression of miRNAs has a neglectable effect on transcriptomes, in contrast to the changes in mRNA expression regulated by the dynamics of hc-siRNA hotspots. Our findings unravel SV-related differential expression of 24-nt hc-siRNAs regulating the expression of certain genes associated with responses to biotic and abiotic stresses as well as differential selection pressures over distinct classes of sRNAs during tomato domestication. Our dataset is also valuable to promote other sRNA-related functional studies on tomato development and domestication.

## Results

### Comprehensive tomato sRNA profiles highlighting expression dynamics as a consequence of domestication

Resulting from domestication, tomatoes underwent substantial changes in plant morphology, yield, fruit flavor and adaptation to adverse environments, reflecting certain levels of adjustments in both genomes and transcriptomes. We selected nine accessions spanning from wild ancestors (*Solanum pimpinellifolium*), semi-domesticated populations (*S. lycopersicum* var. *cerasiforme*) to domesticated tomatoes (*S. lycopersicum* var. *lycopersicum*) for comprehensive transcriptome analyses. These accessions included two SP accessions (BGV006370 collected in Peru and BGV007151 collected in Ecuador), six SLC accessions (BGV005895, BGV007023 and PI 129026 collected in Ecuador; BGV007990 and BGV008189 collected in Peru; BGV008219 collected in Costa Rica), and one SLL accession (BGV007863 collected in Mexico) [1, 3].

For each accession, transcriptome profiles including both sRNA and mRNA profiles were investigated in young leaves, anthesis-stage flowers, and fruits at four different developmental stages (young green, mature green, breaker and red ripe). The four fruit developmental stages have previously been used to analyze the gene regulatory networks in a single accession of domesticated tomato [19, 20]. Therefore, our comprehensive dataset would empower detailed dissection on the gene regulatory networks underlying the fruit ripening process in addition to the transcriptome profiles of leaf and flower.

To ensure sampling at comparable developmental stages of different tomato accessions, we first documented the timing of fruit developments by tracking 10-20 fruits from 3-5 plants of each accession. As shown in **Fig. S1**, the two SP accessions exhibited slightly early ripening whereas SLL ripened one week later. Ripening time in SLC accession varied from as early as SP to as late as SLL. We collected samples at the chosen time points and constructed a total of 162 sRNA-Seq libraries (**Table S1**). The three replicates from young green fruits of BGV007023 did not pass quality check, so we only used the remaining 159 libraries for the subsequent analyses. Principal component analysis (PCA) showed that samples at the same developmental stage were clustered together (**Fig. S2A**). In addition, expression profiles in SLC and SLL accessions were more closely related in comparison to profiles in SP accessions (**Fig. S2B**). Analysis on sRNA size distribution showed that 24-nt siRNAs were the most dominant (**Fig. S2C**). Interestingly, 24-nt siRNAs were slightly more abundant in leaf and young green fruit samples (**Fig. S2C**). To improve the sRNA mapping accuracy, we exploited two high-quality reference genomes (genomes of the domesticated Heinz 1706 and an SP accession LA2093) as detailed in a recent study [5]. It is noteworthy that as expected, reads from SP samples were mapped to the LA2093 genome with a slightly higher rate, while reads from SLC and SLL samples were mapped to the Heinze SL4.0 with a slightly higher rate (**Fig. S2D**). RNA-Seq libraries were also generated from the same samples and described in our previous study [5, 21].

### Hc-siRNA hotspots and structural variants

The 24-nt hc-siRNAs play a major role in RNA-directed DNA methylation, a fundamental mechanism in epigenetic regulation [7, 8]. We reasoned that comparative analyses on hc-siRNA accumulation patterns may provide insights into the dynamic epigenetic changes underlying trait-related gene expression. When analyzing the global hc-siRNA abundance across the 12 tomato chromosomes, we noticed that hc-siRNA abundance displayed a strong genome-wide correlation with the density of SVs (**Fig. 1A**). This observation infers a novel model that epigenetic regulation may be influenced by SVs during crop domestication, which could lead to large scale gene expression changes.

**Figure 1.**
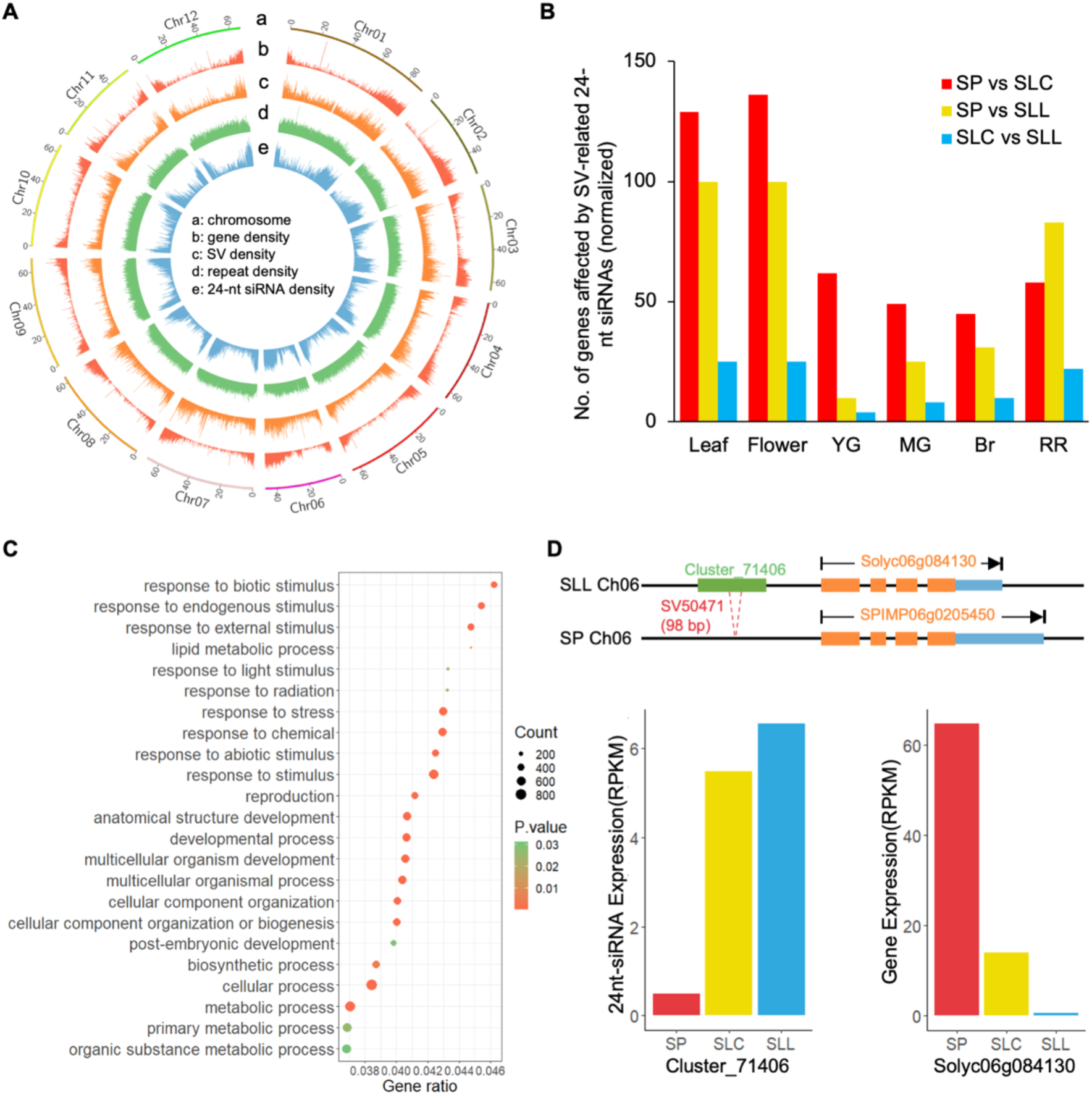
Functional analysis of SV-related 24-nt hc-siRNA regions among wild and cultivated tomatoes. **A**. Circos plot of the densities of 24-nt hc-siRNAs, genes, repeat sequences, and SVs across the tomato genome. **B**. Differentially expressed genes showing a negative correlation with the corresponding SV-related 24-nt hc-siRNA clusters in their promoters in at least one pairwise comparison. YG, young green fruit; MG, mature green fruit; Br, fruit at the breaker stage; RR, red ripe fruit. **C**. GO term enrichment analysis of differentially expressed genes in (**B**). **D**. SV-related 24-nt hc-siRNA (Cluster_71406) enrichment leading to the repression of *Solyc06g084130* expression in SLL (*Solanum lycopersicum* var. *lycopersicum*) and SLC (*S. lycopersicum* var. *cerasiforme*) compared to SP (*S. pimpinellifolium*).

To obtain more evidence in support of this model, we analyzed SV-overlapping hc-siRNA hotspots in promoter regions of protein-coding genes, and identified hc-siRNA and protein-coding gene pairs whose abundances showed simultaneous negative correlations in our sRNA-Seq and RNA-Seq data. For each tissue or development stage, we performed pairwise comparisons between SP and SLL accessions (2 comparisons), between SP and SLC accessions (12 comparisons except 10 for young green fruit stage), and SLC and SLL accessions (6 comparisons except 5 for young green fruit stage), therefore a total of 117 pairwise comparisons for all six tissues/developmental stages. A total of 1,386 protein-coding genes affected by the SV-overlapping hc-siRNA hotspots were identified in at least one comparison. We noticed that most of these genes under the control of this novel epigenetic regulation were expressed in leaf and flower (**Fig 1B**). Gene ontology (GO) term analysis showed that the majority of the protein-coding genes affected by these domestication-associated epigenetic changes were related to pathways in response to biotic and abiotic stresses as well as development and several developmentally related processes (**Fig. 1C**), implying a selection pressure favoring expression changes in genes related to environmental adaptation including selection for domestication traits. To obtain highly confident negative correlations between differentially accumulated 24-nt hc-siRNAs and the cognate protein-coding genes, we focused on the pairs of hc-siRNAs and the negatively correlated protein-coding genes repeated in at least ten pairwise comparisons, which resulted in the identification of 99 protein-coding genes (**Table S2**). The majority of these 99 genes are involved in plant resistance to pathogens. Therefore, SVs appeared to have played a role in shaping the hc-siRNA hotspots across the tomato genome. This regulation over the dynamics of hc-siRNA hotspots has resulted in differential gene expression during tomato domestication, probably under the selection pressure for plant adaptation to different environments and in agricultural settings.

A notable example is a BAX inhibitor-1 (BI-1) family gene (*Solyc06g084130*). BI-1 proteins are conserved in eukaryotic organisms and associated with cell death during host-pathogen interactions [22-24]. In plants, BI-1 proteins regulate the autophagy process [23] and confer plant resistance to various pathogens [23, 25-27]. In particular, BI-1 expression regulates autophagic activity that is critical for *N* gene-mediated resistance to tobacco mosaic virus [23], a major viral pathogen of tomato. The repression of *Solyc06g084130* expression was strongly associated with SV-related 24-nt hc-siRNA differential accumulation from SP to SLC and SLL plants (**Fig. 1D**). Similar repression patterns could be found in multiple genes related to responses to biotic stresses (**Table S2**). Therefore, the repression of *Solyc06g084130* and other genes in response to biotic stresses during tomato domestication may affect tomato resistance traits, which is consistent with the observation that cultivated tomatoes are not well adapted to adverse environments as wild tomatoes.

### Highly dynamic gain/loss of phasiRNA generating loci during domestication

PhasiRNAs are known regulators of gene expression in plants [28] and exhibit expression dynamics in response to environmental cues [13]. Current models suggest that phasiRNAs serve as negative regulators to modulate the expression of their parental transcripts [13, 28]. Although there is rapid progress in uncovering an enormous amount of phasiRNA-generating loci in various plants [13, 28] and unraveling their functions in plant development [29-31], a detailed analysis of phasiRNA dynamics during crop domestication has not been conducted. Using a previously established algorithm [32], we analyzed phasiRNA-generating loci across all nine accessions and identified 290 PHASs mapped to the Heinz 1706 genome (SL4.0) and 286 mapped to the LA2093 genome, among which 77 were uniquely mapped to SL4.0 and 73 uniquely mapped to the LA2093 genome (**Tables S3-5**). In general, PHASs were mainly mapped to protein-coding genes with diverse functions as shown in the GO term analysis (**Fig. S3A**), akin to our previous findings [13]. Interestingly, we found 34 SV-overlapping PHASs mapped to SL4.0 and 39 mapped to the LA2093 genome, among which 12 were uniquely mapped to SL4.0 and 17 uniquely mapped to the LA2093 genome (**Table S6**). SV-related PHASs were distributed across all chromosomes except chromosome 4 and 7, and there were numerous SV-related PHASs residing proximal to the terminal region of the long arm of chromosome 11 (**Fig. 2A**). SVs markedly contributed to the changes in PHASs among accessions in different tomato groups (**Fig. S3B**), and these PHASs were overwhelmingly mapped to disease resistance genes (**Fig. S3C**). This observation hints the possibility that selection pressure favors the emergence of phasiRNAs regulating the expression of disease resistance genes in balancing growth and pathogen defense.

**Figure 2.**
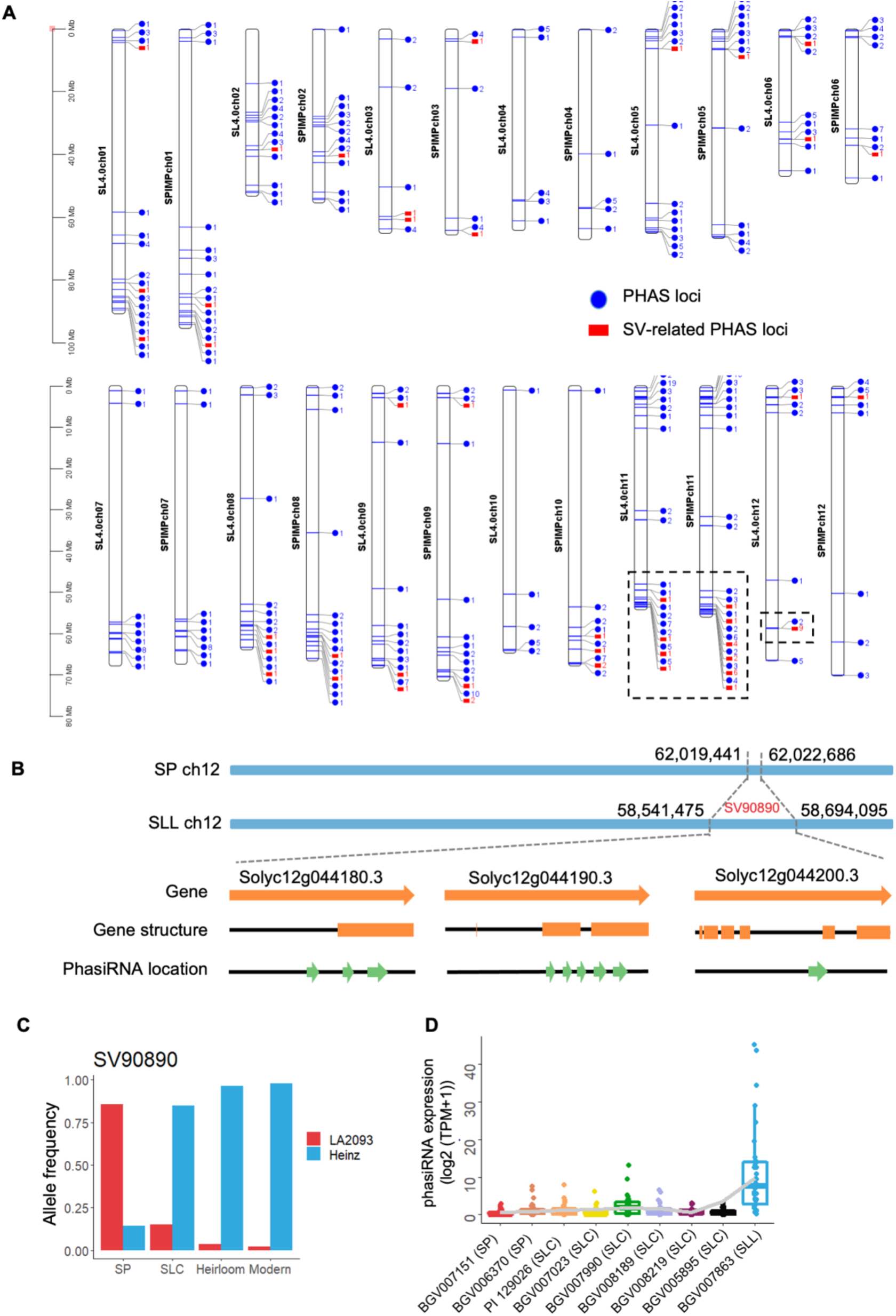
SV-related PHASs. **A**. Distribution of PHAS loci and SV-related PHAD loci across the 12 tomato chromosomes. The number depicts the distinct PHASs at each locus. **B**. SV90890 causes gain/loss of PHASs and protein-coding genes. **C**. Allele frequency of SV90890 in different tomato groups. **D**. Abundance of phasiRNAs in the genome region of SLL Heinz 1706 containing SV90890 in the nine tomato accessions. For each box plot, the lower and upper bounds of the box indicate the first and third quartiles, respectively, and the center line indicates the median. The whisker represents 1.5× interquartile range of the lower or upper quartile.

One cluster of disease resistance genes resided in a region where an insertion in the Heinz 1706 genome expanded the PHASs on chromosome 12 (**Fig. 2B**). All three genes in this inserted region are involved in tomato resistance to bacterial and oomycete pathogens [33]. As shown in **Fig. 2C**, the frequency of the insertion associated with this gene cluster drastically increased in domesticated tomato to nearly 100%. Productions of phasiRNAs associated with this SV were also highly elevated in the SLL accession (**Fig. 2D**).

### A large portion of miRNAs exhibit highly dynamic expression patterns

The functions of many miRNAs in tomato growth and fruit development have been well studied in single accession analyses [20, 34, 35]. However, detailed analyses on miRNA expression profiles are lacking to infer the dynamics of miRNA-based gene regulatory network during crop domestication. To this end, we annotated all the miRNAs in our sRNA-Seq dataset based on recently revised criteria [36]. We identified 122 miRNA genes mapped to SL4.0 and 126 mapped to the LA2093 genome. Both sets included 72 previously reported tomato miRNAs [20, 37, 38]. There were 116 miRNA genes mapped to both SL4.0 and the LA2093 genome, while there were six and ten miRNA genes specifically mapped to SL4.0 and the LA2093 genome, respectively (**Table S7**). Based on the mapping results, we summarized the mature miRNAs and miRNA*s as well as the processing variants from the miRNA precursors in **Table S8-9**.

A close look at the mature miRNAs showed that the majority of known miRNAs were 21-nt in length with “U” as the first nucleotide (**Fig. S4**), in line with previous observations [12]. Notably, the majority of novel miRNAs were 20-nt in length with “A” as the first nucleotide (**Fig. S4**), which implied that those miRNAs could lack regulatory functions. To our surprise, we found that a large portion of miRNAs exhibited highly variable expression patterns among accessions. For example, as shown in **Table S10**, the two SP accessions each has ∼120 miRNAs that displayed 1.5-fold changes in expression with an adjusted P value below 0.05 compared with SLC and SLL accessions.

To better describe this dynamic, we plotted miRNA expression profiles across the nine tomato accessions and categorized the patterns into multiple groups. As shown in **Fig. 3**, there were eight groups with each having more than ten distinct miRNAs. While the majority of the miRNAs exhibited dynamic expression profiles in SLC accessions, we found that those in groups 1 and 7 exhibited an overall decrease while those in groups 5, 6, and 8 exhibited an overall increase in expression over the course of tomato’s recent evolution. The presence of eight distinct groups also reflected the fluctuating expression patterns of miRNAs among different tomato accessions. However, we did not observe the corresponding changes in the expression of their targets, which will be further analyzed below.

**Figure 3.**
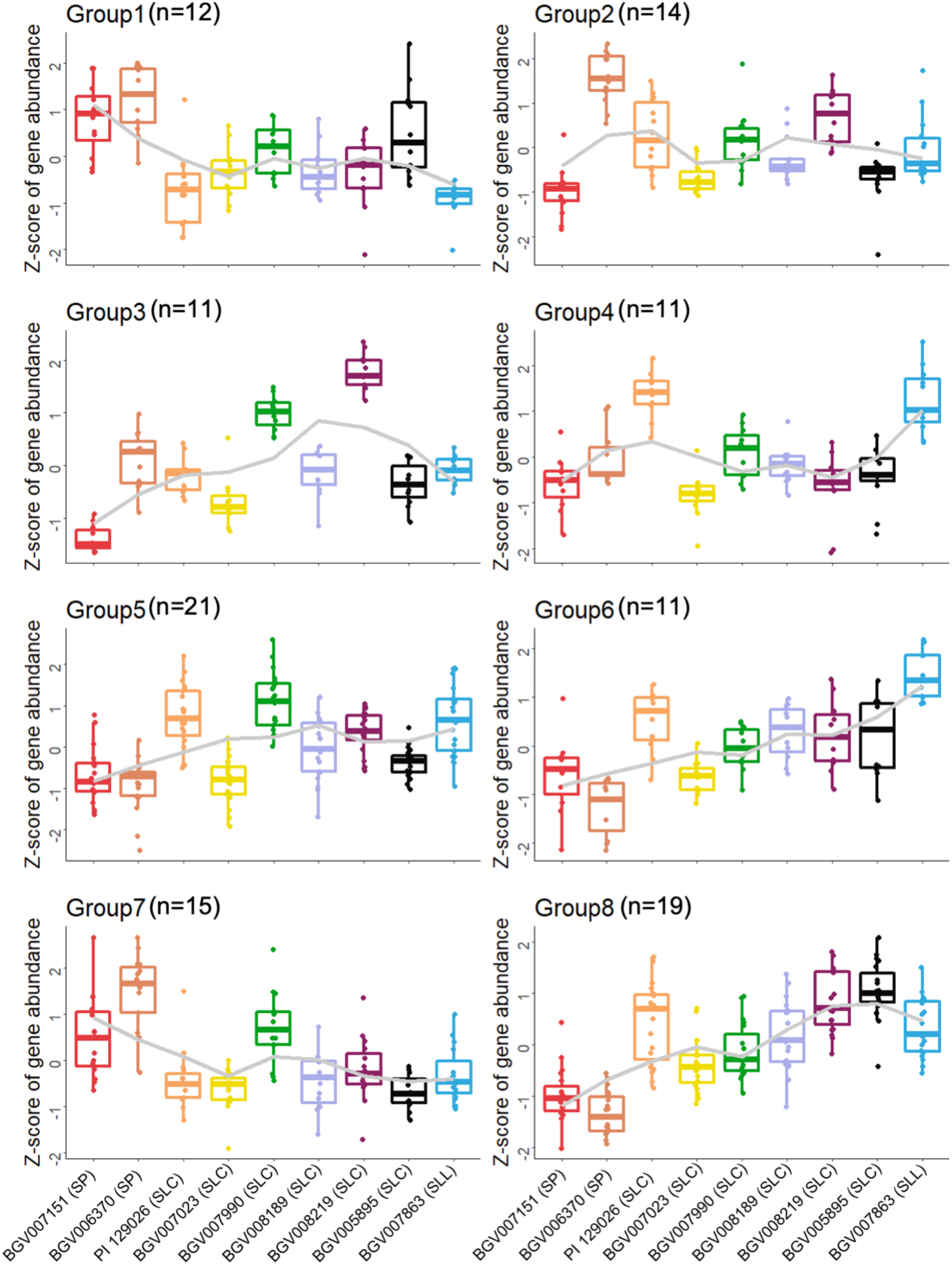
Distinct expression patterns of miRNAs. For each box plot, the lower and upper bounds of the box indicate the first and third quartiles, respectively, and the center line indicates the median. The whisker represents 1.5× interquartile range of the lower or upper quartile.

### miRNAs and structure variants

Despite the notion that most miRNA gene families had the same number of members mapped to SL4.0 and the LA2093 genome, some underwent deletion or duplication events that changed the number of members in each family. For example, there were two miR10535 genes in the LA2093 reference genome but only one copy in Heinz 1706 SL4.0 (**Fig. 4A**). Synteny analysis showed that the loss of one miR10535 gene copy could be attributed to a deletion that had occurred during tomato domestication (**Fig. 4A**).

**Figure 4.**
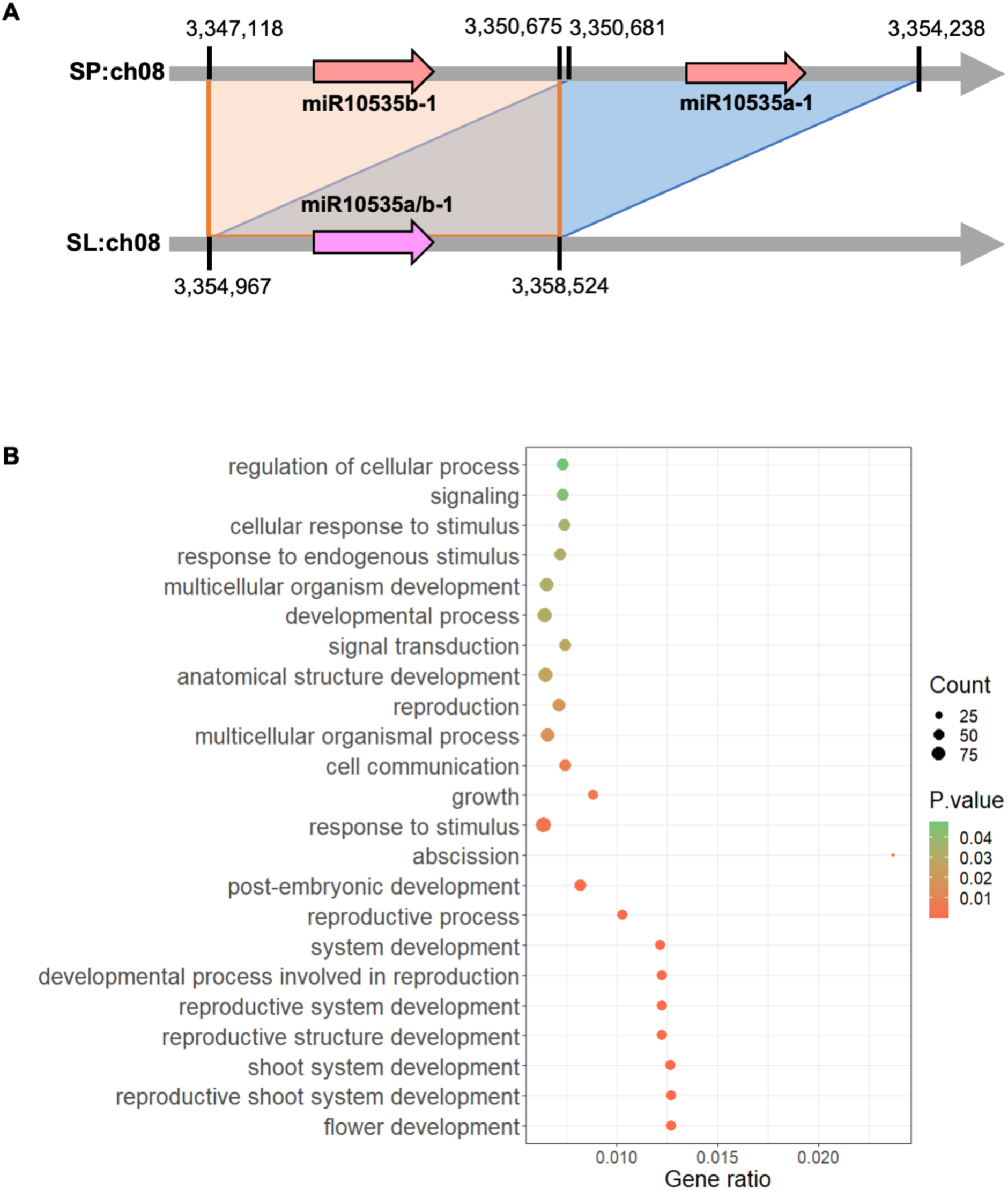
MiRNA genes overlapped with SVs. **A**. Synteny diagram of miR10535 gene(s) in SP and SLL reference genomes. **B**. Enriched GO terms of predicted target genes of SV-related miRNAs.

Since a genome deletion caused the loss of one copy of miR10535 gene, we reasoned that SVs might play important roles in determining the presence and the expression levels of miRNA genes. To this end, we identified 25 miRNA genes associated with 32 SVs, including 19 conserved and six novel miRNAs. Twenty-one out of 32 SVs were mapped to promoters of miRNA genes, seven were mapped to miRNA gene bodies, and four were mapped to both promoters and gene bodies of miRNAs (**Table S11**). Notably, most of these miRNAs do not have empirically confirmed targets according to previous degradome studies in tomato [20, 39-41]. GO term analysis showed that the computationally predicted target genes of SV-overlapping miRNAs were mainly involved in the development/growth and responses to environmental stimuli (**Fig. 4B**).

SVs in promoter regions may affect miRNA expression. For example, a 17-bp deletion (SV49979) in the promoter of the miR172a/b-2 gene was observed in genomes of most heirloom and modern tomatoes (**Fig. 5A**). It is worth noting that the allele frequency of SV49949 significantly changed during domestication (**Fig. 5B**). Correspondingly, miR172a/b-2 had distinct expression profiles during fruit development among the nine tomato accessions and showed an overall increased expression pattern in the SLC and SLL accessions (**Fig. 5C**). Due to the low abundance of miR172a/b-2 compared to the total miR172 abundance (**Tables S8 and S9**), this expression change in miR172a/b-2 did not significantly affect the expression of any miR172 targets.

**Figure 5.**
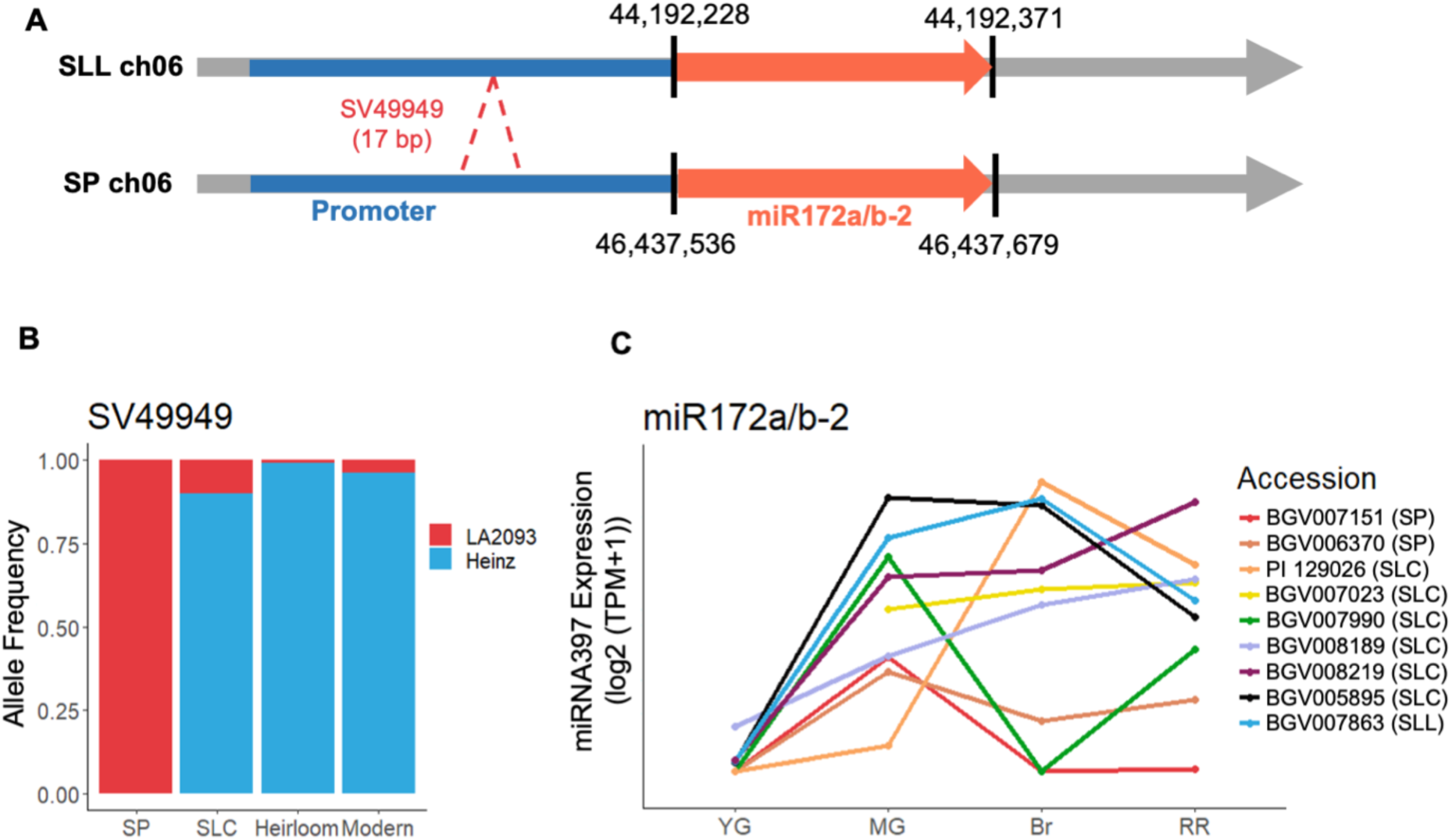
SV affects miRNA expression. **A**. Diagram showing SV49949 (a 17-bp indel) in the promoter of miR172a/b-2 gene. **B**. Allele frequency of SV49949 in different tomato groups. **C**. Expression profiles of miR172a/b-2 in different tomato accessions. YG, young green fruits. MG, mature green fruits. Br, fruits at the breaker stage. RR, red ripe fruits.

### Imprecisely processed miRNAs are differently selected

It is known that some miRNA precursors tend to generate a population of miRNA-miRNA* pairs due to the imprecise processing by DCL1 [42]. The imprecise processing may affect miRNA function that heavily relies on sequence complementarity between miRNAs and their targets [43]. We found eight out of 122 SLL miRNA genes and six out of 126 SP miRNA genes that generated more than six variants in our dataset, among which six were shared in both SLL and SP reference genomes (**Table S12**). The miR397 gene, a conserved miRNA across different plant lineages, had six miR397-3P and seven miR397-5P variants in tomato (**Fig. 6A**). Notably, we observed a shift in the most abundant product of the miR397 precursor: from only miR397-3P that was expressed in SP accessions transitioning to miR397-5P (the conserved mature miR397 in plants) that became more prevalent in most SLC and SLL accessions (**Fig. 6B**). Over-expression of miR397-5P can enhance tomato response to drought [44], suggesting the beneficial function of miR397-5P in SLC and SLL accessions in adaptation to adverse environments. Nevertheless, the target of miR397-5P remains unclear in spite of extensive efforts of degradome analysis [20, 39-41].

**Figure 6.**
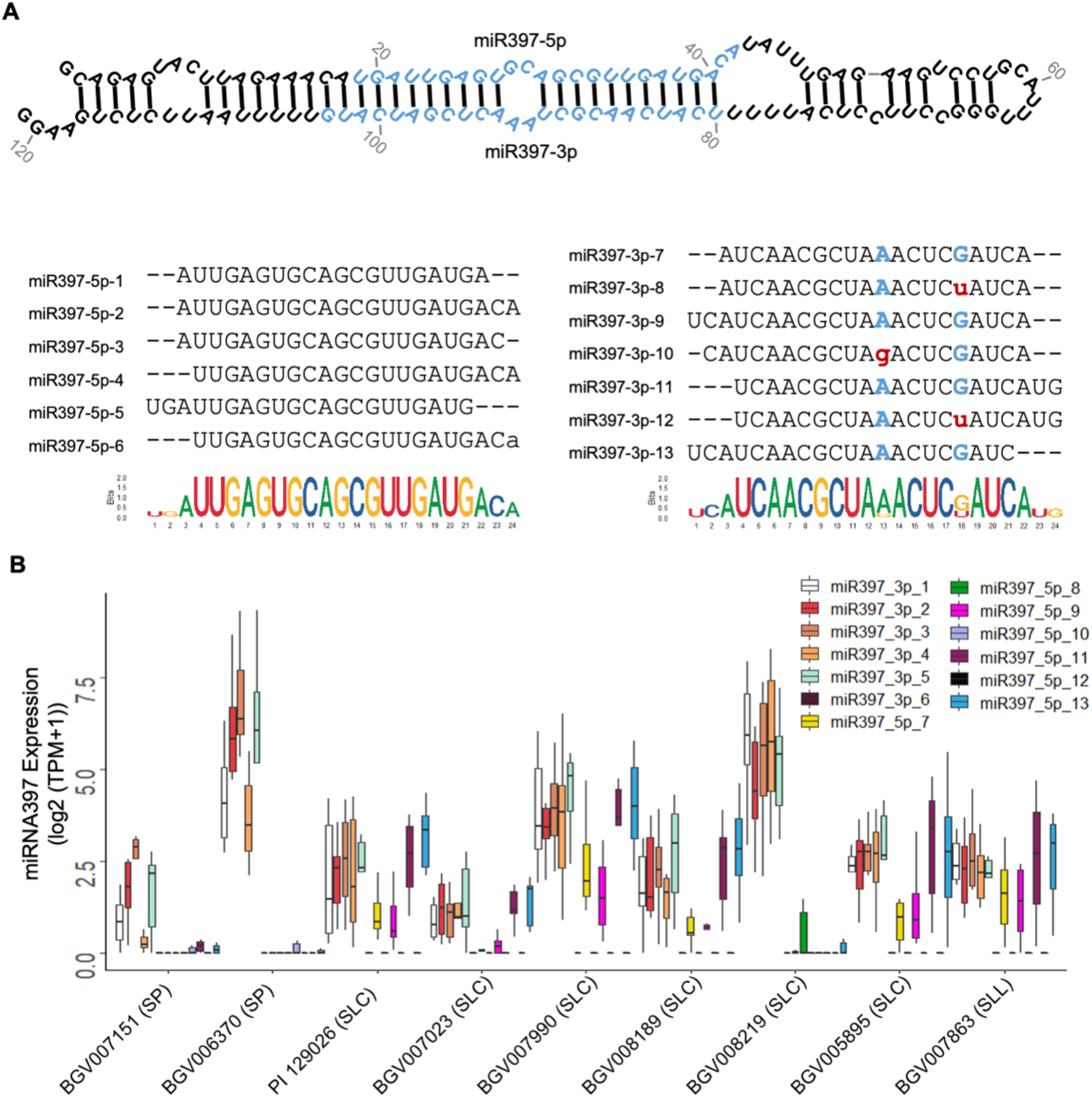
Changes in miR397 expression during tomato domestication. **A**. Diagram showing the six 5P and seven 3P processing variants from the miR397 precursor. **B**. Abundance of each miR397 variant in different tomato accessions.

We also observed a position shift of miRNA:miRNA* in the precursor of miR9472. The miRNA:miRNA* duplex was closer to the terminal loop region in SP and SLC accessions but resided at a more distant region away from the terminal loop in SLL (**Fig. S5**). This shift was unlikely caused by any changes in recognition by DCL1 as the precursor sequences remained the same. Notably, we found seven such examples as listed in **Table S13**. This observation indicates that DCL1 recognition on miRNA precursors is likely flexible in plants, and the selection pressure plays a role in determining the expression of the final miRNA:miRNA* duplexes.

## Discussion

Small RNAs are critical regulators of gene expression underlying tomato growth and responses to environmental cues [20]. sRNA abundance is known to directly impact their functions. To gain a better understanding of sRNA dynamic expression during tomato domestication, we generated a comprehensive sRNA dataset using nine representative tomato accessions spanning from the wild SP progenitors, intermediate SLC accessions, and one domesticated accession, and covering samples from leaf, flower, and fruits at four critical developmental stages. Our high-quality dataset fulfills the immediate needs for high-resolution comparative analyses on sRNA expression and inferring their functions in wild, semi-domesticated, and domesticated tomato plants, as well as establishes a foundation for future exploration of sRNA functions.

Our dataset clearly demonstrates that all three classes of sRNAs (hc-siRNAs, phasiRNAs and miRNAs) have significant changes in expression during tomato’s recent evolution. Notably, we found that SVs are an important driving force underlying the dynamic expression of these sRNAs. Those SVs, particularly deletions and insertions, probably have a direct impact on the production of all three major types of sRNAs. For example, deletions or insertions result in the differential accumulation of hc-siRNAs in gene promoter regions and the gain or loss of PHASs. SVs also contribute to the birth and death of miRNAs in domesticated tomatoes, as evidenced by the deletion of miR10535 and miR482f. When SVs reside in promoter regions of miRNA genes, they may influence miRNA expression, as evidenced in miR172a/b-2. In addition to SVs, we also notice that imprecise processing of miRNA/miRNA* duplexes can lead to the dynamic expression of miRNAs during tomato domestication.

Tomato has a complex history of domestication, selection and breeding [1-3, 18]. Tomato domestication before cultivation possibly has a selection pressure on plant adaptation to new environments and developmental processes. Domestication and re-domestication processes possibly have posed a selection pressure on tomato flavor and yield. The modern breeding processes possibly have posed a selection pressure on disease resistance. Interestingly, selection pressures appear to have distinct impacts on different classes of sRNAs. The SV-related dynamic expression of 24-nt hc-siRNAs and phasiRNAs predominantly impacts the expression of genes related to stress responses and growth, implying that the selection pressure favors the regulation of those trait-associated genes through hc-siRNA and phasiRNA pathways. In contrast, the conserved miRNAs play a major role in plant development mostly through regulating transcription factors [14, 15]. Some of those well-established regulations are conserved in most land plants [45-47]. Therefore, the selection pressure is unlikely to favor changes in such critical regulations related to plant growth in a relatively short timeframe during domestication. By contrast, many less conserved miRNAs, mostly expressed at low levels and/or having no to very few confirmed targets, exhibit highly dynamic expression during tomato domestication, reflecting little selection pressure on those miRNAs [14, 15].

Our dataset serves as a foundation for future studies on sRNA function associated with tomato growth, domestication, and beyond. Recent progress has demonstrated that miRNA gene families may exert functions through developmentally-regulated expression of specific members [48, 49]. Our dataset can help mining miRNA family members with developmentally-regulated expression patterns. For example, miR390b, with a U20A substitution at position 20 in comparison to miR390a, was specifically expressed in flowers (**Fig. S6**). The flower-specific expression of miR390b tripled the amount of total miR390 in flowers, which markedly promoted production of phasiRNAs from the *TAS3* locus as well as specifically suppressed the expression of *ARF3* and *ARF4* in flowers (**Fig. S6**).

## Methods

### Plant materials and RNA isolation

Tomato plants were grown in a greenhouse at 25 °C and with a 16/8 hr light/dark cycle at Ohio State University (Columbus, OH). For each accession, young leaves, anthesis-stage flowers, and fruits at four different developmental stages (young green, mature green, breaker and red ripe) were collected with three biological replicates. Total RNAs from tomato samples were isolated and fractioned to >200 nt and <200 nt populations using the RNAzol RT reagent (Sigma-Aldrich, St. Louis, MO). sRNA species were further purified using the miRVana miRNA isolation kit (Thermo Fishier Scientific, Grand Island, NY) following the manufacturer’s instructions. mRNA populations were further purified using the Magnetic mRNA isolation kit (NEB, Ipswich, MA).

### Library construction and sequencing

sRNA libraries were constructed following the established protocol [50]. Briefly, 18-30 nt sRNA populations purified on 15% (w/v) polyacrylamide/8M urea gel were ligated with 3’- and 5’-adapters. sRNA populations with adapters were reverse transcribed, PCR amplified, and then purified from the 8% native PAGE gel. Strand-specific RNA-Seq libraries were constructed using the protocol described before [51]. All the constructed libraries were analyzed and quantified by Bioanalyzer and sequenced on an Illumina HiSeq 2500 system.

### sRNA sequence processing

sRNA reads were processed to remove adaptors using the sRNA cleaning script provided in the VirusDetect package [52]. The trimmed sRNA reads shorter than 15 nt were discarded. The resulting sRNA reads were further cleaned by removing those that perfectly matched to the sequences of tRNAs, snoRNAs, snRNAs (collected from GenBank) or rRNAs [53] using Bowtie [54]. Raw counts for each unique sRNAs were derived and normalized into TPM (transcripts per million).

### Identification of miRNAs and differential expression analysis

MiRNAs were identified using ShortStack [55] from each of the 159 samples, and a series of filtering was applied to obtain high-confidence miRNAs. Briefly, the cleaned sRNA reads were mapped to wild (LA2093) and cultivated tomato (Heinz 1706, SL4.0 and ITAG 4.1) genomes separately, using ShortStack [55] with the parameter ‘mmap’ set to ‘u’. Mature miRNAs and corresponding pre-miRNAs were then identified by ShortStack [55]. The identified miRNAs from these samples were collapsed if they were mapped to the exact locations in the genome. The collapsed miRNAs that existed in at least three samples and expressed at more than 10 TPM were considered as high-confidence miRNAs, which were compared with miRBase [56] to identify conserved miRNAs, whereas miRNAs that showed no matches in the miRBase were considered as novel miRNAs.

Raw counts of the identified miRNAs were processed using DESeq2 [57] to identify differentially expressed miRNAs among accessions. MiRNAs with adjusted p values < 0.05 were considered as differentially expressed. Differentially expressed miRNAs were further clustered into groups according to their expression patterns using the DEGreport program [58]. Target genes of differentially expressed miRNAs were predicted using the TargetFinder program [59]. GO enrichment analysis was performed on the target genes using GO::TermFinder [60].

### Identification of candidate PHAS loci

We used the previously described methods to identify PHAS loci [13]. In brief, the cleaned sRNA reads were mapped to the wild (LA2093) and cultivated tomato (Heinz 1706, SL4.0 and ITAG 4.1) reference genomes using Bowtie [54] allowing no mismatch and no more than six hits. The reference sequences were then scanned with a sliding window of 189 bp (nine 21-nt phase registers). A positive window was considered to contain no less than 10 unique sRNAs, with more than half of unique sRNAs being 21 nt in length and with no less than three 21-nt unique sRNAs falling into the phase registers. Windows were combined if 1) they shared the same phase registers and 2) fell into the same gene loci. P values and phasing scores for positive windows were calculated following the methods described previously [32, 61]. The sequences of PHASs and the flanking regions of 200 bp were retrieved and compared between wild and cultivated tomato reference genomes using the BLAST program [62]. The BLAST results were then processed to categorize candidate PHAS loci into three groups: PHAS loci shared by wild and cultivated tomatoes, specific to wild tomato, and specific to cultivated tomato.

### Identification and analysis of 24-nt hc-siRNA hotspots

Cleaned sRNA reads from all 159 samples were fed into ShortStack [55] to identify siRNA hotspot regions that were defined by continuously covered sRNAs. The expression of 24-nt siRNA was calculated by counting the number of 24-nt siRNA reads mapped to the corresponding regions. Only a region with no less than ten mapped 24-nt siRNA reads was considered as a 24-nt siRNA hotspot. The 24-nt siRNA expression was normalized to number of reads per kilobase of region per million mapped reads (RPKM), based on all mapped reads.

To identify SV-related 24-nt siRNAs, pairwise comparisons were firstly applied between stages or between tissues, and statistical analysis was performed using DESeq2 [57]. Only regions with adjust P-values < 0.05 and fold change ≥ 2 were considered as significantly changed hotspots. The significantly changed protein-coding genes and corresponding changed 24-nt siRNA hotspots in promoter regions were treated as genes pairs involved in epigenetic regulation. To further exclude potential false positive candidates, we only kept the pairs of 24-nt hotspots and the corresponding protein-coding genes with a negative correlation in expression that occurred in at least 10 samples. Previously reported SVs [5] that overlapped with significantly changed 24-nt siRNA hotspots were then identified. The identified SVs were further filtered to keep those in promoter regions of protein-coding genes that exhibited an opposite expression pattern compared to the changes in abundance of the corresponding 24-nt siRNA hotspots.

### RNA-Seq read processing and differential expression

Single-end RNA-Seq reads were processed to remove adapters as well as low-quality bases using Trimmomatic [63], and the trimmed reads shorter than 80 bp were discarded. The remaining high-quality reads were subjected to rRNA sequence removal by aligning them to an rRNA database [53] using Bowtie [54] allowing up to three mismatches. The cleaned RNA-Seq reads were aligned to the cultivated tomato (Heinz 1706, SL4.0) reference genome using STAR [64] allowing up to two mismatches. Gene expression was measured by counting the number of reads mapped to gene regions (ITAG4.1), and then normalized to the number of reads per kilobase of exon per million mapped reads (RPKM). Differential expression analysis was performed using DESeq2 [57]. To obtain a global comparison among all samples, in particular to identify differentially expressed genes in specific accessions or developmental stages, we followed a previously described linear factorial modeling [65]. We also performed pairwise comparisons to identify differentially expressed genes between stages for different accessions. Genes with adjusted P values <0.05 and fold changes no less than two were considered differentially expressed.

## Accession numbers

Raw sRNA and RNA-Seq reads have been deposited in the NCBI SRA under the accession number SRP135718.

## Acknowledgements

This work was supported by grants from the US National Science Foundation (IOS-1564366 to EvdK and YW and IOS-1855585 to ZF), the Urban Agriculture Characteristic Platform Construction of Beijing University of Agriculture (5076016177/003 to YZ). This work is dedicated to the late Prof. Biao Ding (Ohio State University), who participated at the inception stage of this study.

## Author contributions

YZ, EvdK, YW, and ZF conceived the idea. YZ, YW, and ZF supervised the study. SM and YW performed the plant analyses and sampling. YW constructed sRNA-Seq libraries. YQ, YZ, HNS, XW, XZ, YW, and ZF analyzed the data. YZ and YW summarized the results. YZ, EvdK, YW, and ZF wrote and revised the manuscript. All authors have read and approved the final manuscript.

