## Supplementary material for "Large-scale comparative small RNA analyses reveal genomic structural variants in driving expression dynamics and differential selection pressures on distinct small RNA classes during tomato domestication": Figure S1

**A**

| Accession | Origin | Group | # of fruits | Days post anthesis (average) | | | |
|---|---|---|---|---|---|---|---|
| | | | | Young green | Mature green | Breaker | Red ripe |
| BGV007151 | Ecuador | SP | 20 | 5 | 24.2 | 28.4 | 32.3 |
| BGV006370 | Peru | SP | 12 | 5 | 25.6 | 31 | 35.8 |
| PI 129026 | Ecuador | SLC | 20 | 5 | 29.6 | 32.2 | 37.5 |
| BGV007023 | Ecuador | SLC | 28 | 5 | 27.8 | 30.5 | 34.1 |
| BGV007990 | Peru | SLC | 21 | 7 | 32.6 | 38.1 | 43.2 |
| BGV008189 | Peru | SLC | 22 | 5 | 28 | 33.4 | 37.5 |
| BGV008219 | Mexico | SLC | 32 | 5 | 26.6 | 30.4 | 35.7 |
| BGV005895 | Mexico | SLC | 25 | 5 | 30.4 | 35.4 | 40.9 |
| BGV007863 | Mexico | SLL | 23 | 7 | 32.8 | 37.8 | 42.4 |

**B**

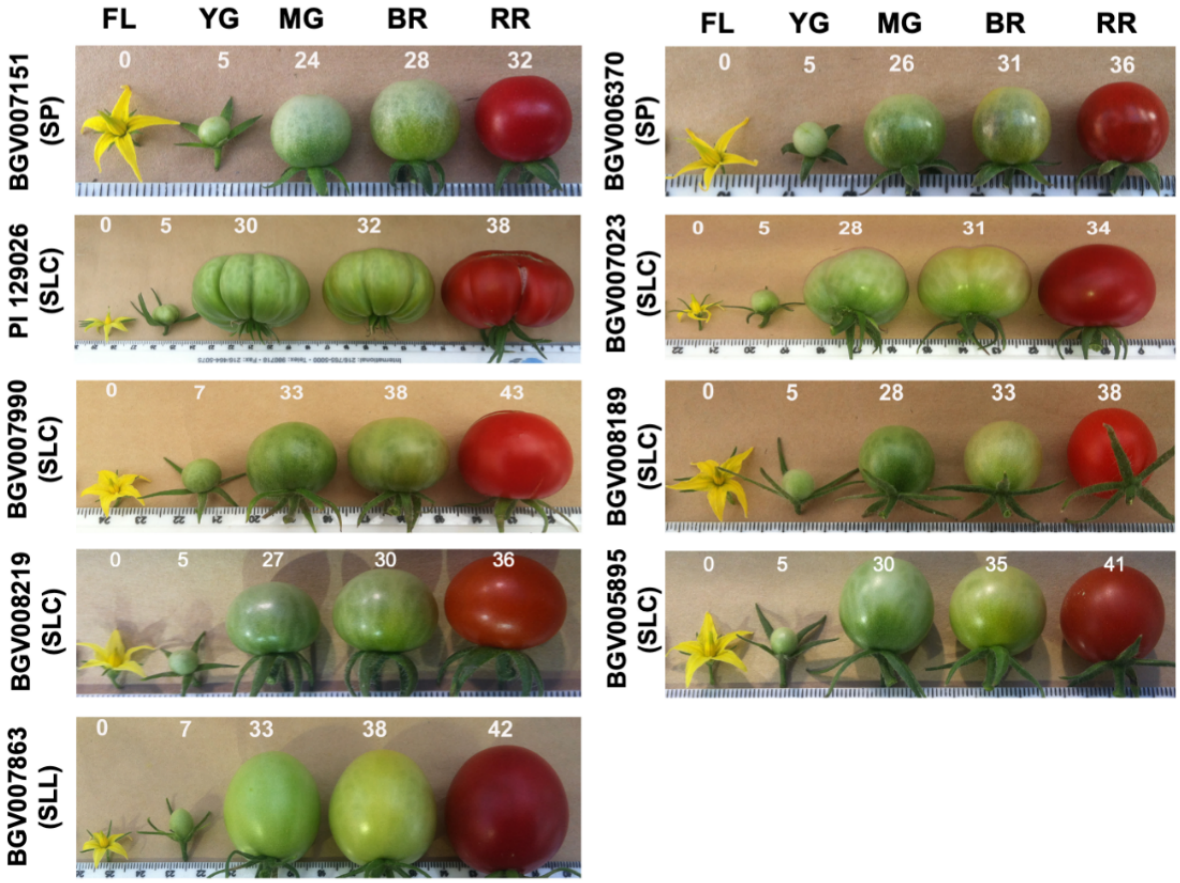


**Figure S1.** Development timeline of the nine tomato accessions investigated in this study. **A.** Summary of fruit development and ripening timelines. **B.** Sample morphologies. FL, flower. YG, young green fruits. MG, mature green fruits. Br, fruits at the breaker stage. RR, red ripe fruits.
