## Supplementary material for "Large-scale comparative small RNA analyses reveal genomic structural variants in driving expression dynamics and differential selection pressures on distinct small RNA classes during tomato domestication": Figure S2

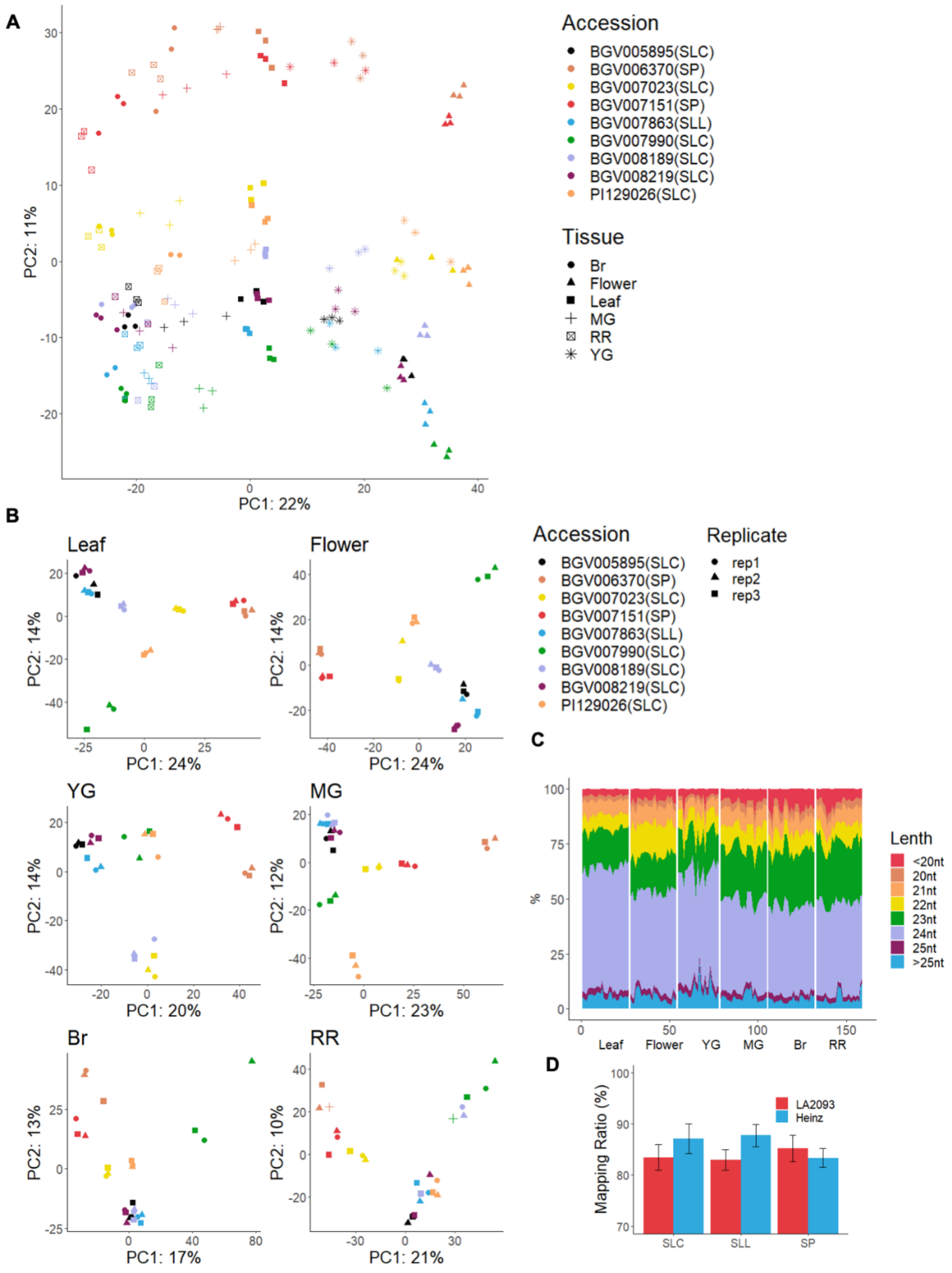


**Figure S2.** sRNA sequencing data. **A.** Principal component analysis (PCA) of all samples. **B.** PCA of samples at each tissue or developmental stage. **C.** Size distribution of sRNAs. **D.** Mapping ratio of sRNAs to Heinz 1706 and LA2093 genomes. YG, young green fruit; MG, mature green fruit; Br, fruit at the breaker stage; RR, red ripe fruit.
