## Supplementary material for "Large-scale comparative small RNA analyses reveal genomic structural variants in driving expression dynamics and differential selection pressures on distinct small RNA classes during tomato domestication": Figure S3

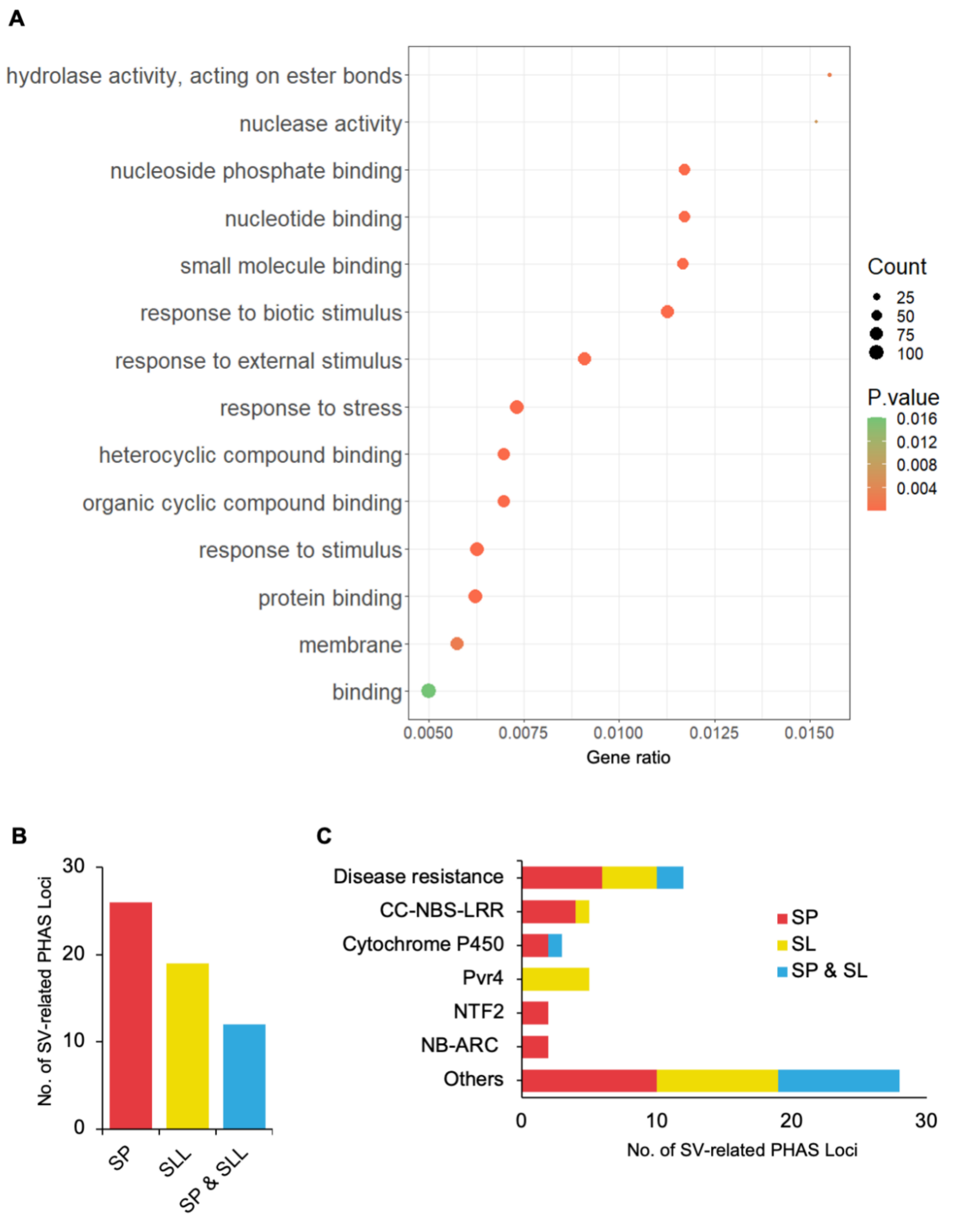


**Figure S3.** Characteristics of PHAS loci. **A.** Enriched GO terms of protein-coding genes that generate phasiRNAs. **B.** SV-related PHASs in wild (SP) and cultivated (SLL) tomatoes. **C.** Functional classification of genes overlapped with SV-related PHASs.
