## Supplementary material for "Large-scale comparative small RNA analyses reveal genomic structural variants in driving expression dynamics and differential selection pressures on distinct small RNA classes during tomato domestication": Figure S4

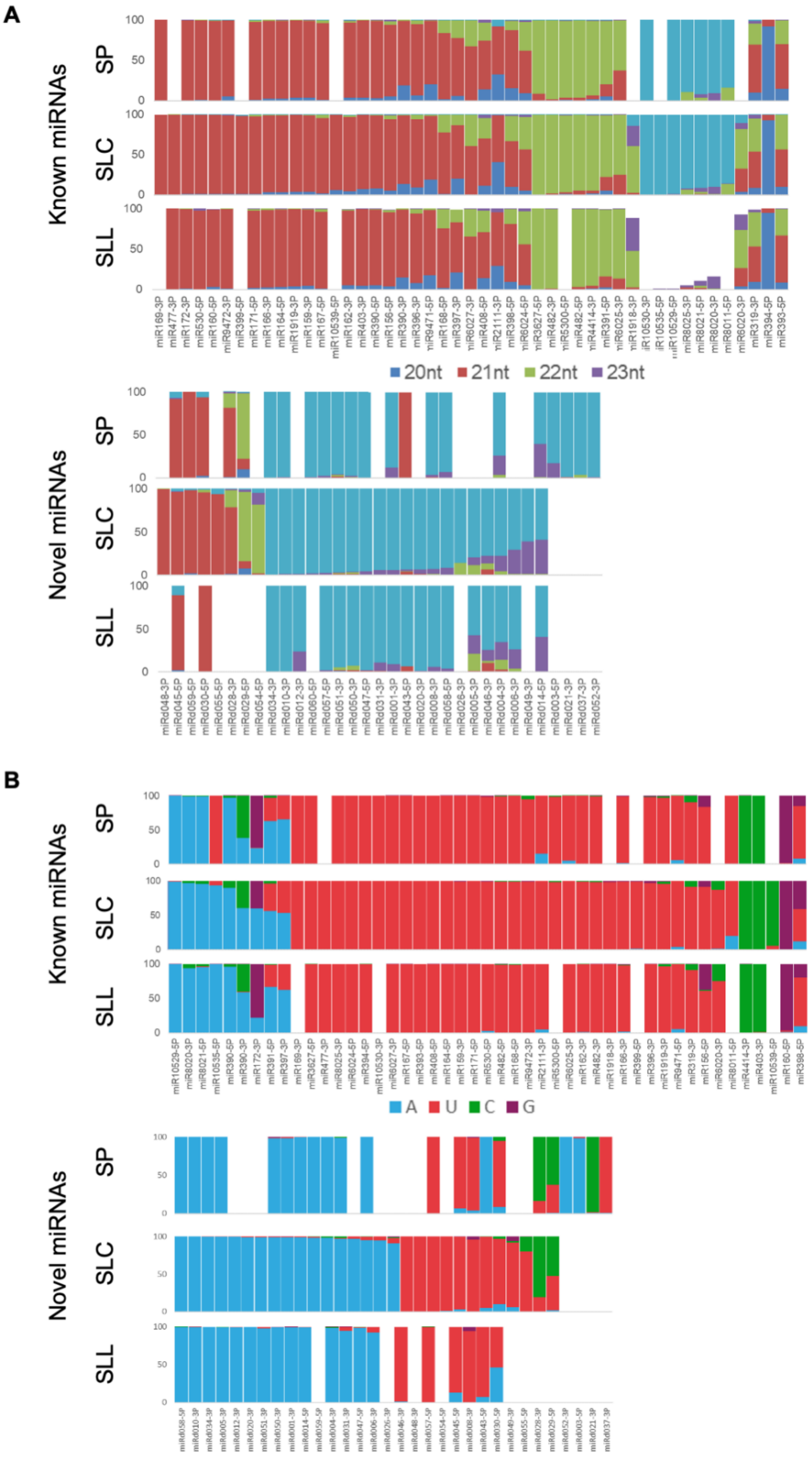


**Figure S4.** Characteristics of mature miRNAs. **A.** Proportion of mature miRNAs with different sizes in different tomato groups. **B.** Composition of the first nucleotide in mature miRNAs.
