## Supplementary material for "Large-scale comparative small RNA analyses reveal genomic structural variants in driving expression dynamics and differential selection pressures on distinct small RNA classes during tomato domestication": Figure S5

**A**

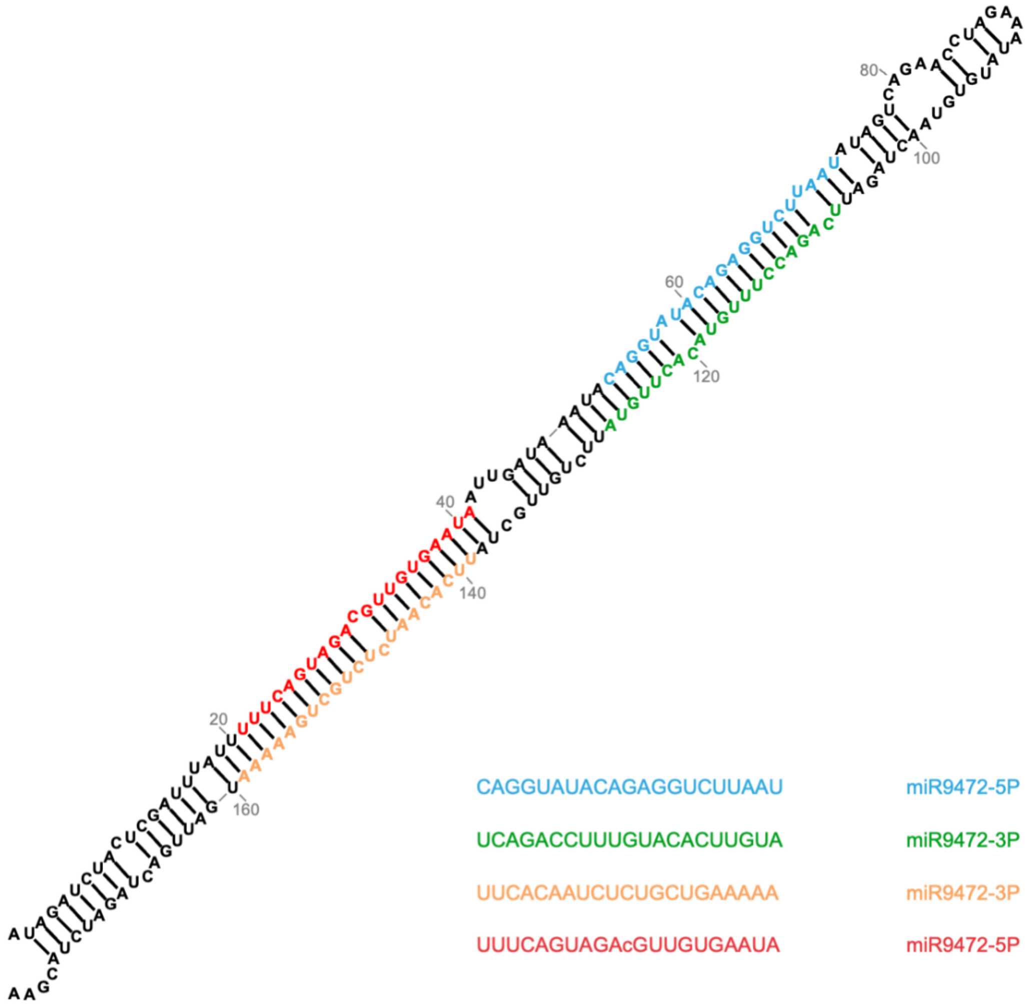


**B**

| pre-miRNA | | | | miRNA | | | No. miRNA (Accession) | | | | | | | | |
|---|---|---|---|---|---|---|---|---|---|---|---|---|---|---|---|
| Chromosome | Start | End | strand | sequence | family | Length | BGV006370 (SP) | BGV007151 (SP) | BGV005895 (SLC) | BGV007023 (SLC) | BGV007990 (SLC) | BGV008189 (SLC) | BGV008219 (SLC) | PI009026 (SLC) | BGV007863 (SLL) |
| SL4.0ch08 | 57078559 | 57078737 | + | CAGGUAUACAGAGGUCUUAAU | Sly-miR9472-5P | 21 | 0 | 1 | 1 | 0 | 1 | 1 | 0 | 0 | 0 |
| SL4.0ch08 | 57078559 | 57078737 | + | UCAGACCUUUGUACACUUGUA | Sly-miR9472-3P | 21 | 0 | 1 | 1 | 0 | 1 | 1 | 0 | 0 | 0 |
| SL4.0ch08 | 57078559 | 57078737 | + | UUCACAAUCUCUGCUGAAAAA | Sly-miR9472-3P | 21 | 0 | 0 | 0 | 0 | 0 | 0 | 0 | 0 | 2 |
| SL4.0ch08 | 57078559 | 57078737 | + | UUUCAGUAGACGUUGUGAAUA | Sly-miR9472-5P | 21 | 0 | 0 | 0 | 0 | 0 | 0 | 0 | 0 | 2 |

**Figure S5.** Shifted miRNA-miRNA* pair from the miR9472 precursor. **A.** Structure of the miR9472 precursor with two alternative miRNA:miRNA* pairs highlighted with different colors. **B.** Presence of the two miRNA:miRNA* pairs of miR9472 in the nine tomato accessions.
