## Supplementary figures and images for "Large-scale comparative small RNA analyses reveal genomic structural variants in driving expression dynamics and differential selection pressures on distinct small RNA classes during tomato domestication"

### Figure S6

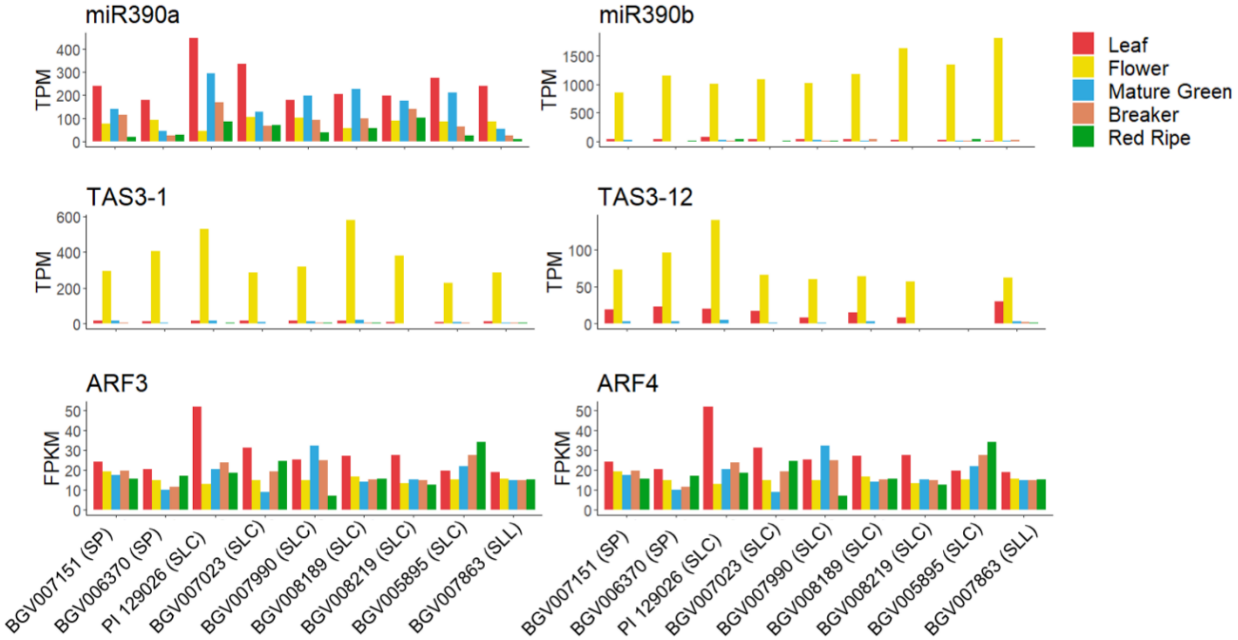


**Figure S6.** Expression profiles of the miR390-TAS3-ARF3/ARF4 cascade in the nine tomato accessions.
